# Simulated colonic fluid replicates the *in vivo* growth capabilities of *Citrobacter rodentium* mutants and highlights a critical role for the Cpx envelope stress response in mediating stressors encountered in the gastrointestinal tract

**DOI:** 10.1101/2021.12.13.472529

**Authors:** Ashley Gilliland, Christina Gavino, Samantha Gruenheid, Tracy Raivio

**Affiliations:** Department of Biological Sciences, University of Alberta, Edmonton, Alberta, Canada; Department of Microbiology and Immunology, McGill University, Montreal, Quebec, Canada

## Abstract

*Citrobacter rodentium* is an attaching and effacing (A/E) pathogen used as a model for enteropathogenic and enterohemorrhagic *Escherichia coli* infections in mice. While in the host, *C. rodentium* must adapt to stresses in the gastrointestinal tract such as antimicrobial peptides, pH changes, and bile salts. The Cpx envelope stress response (ESR) is a two-component system used by some bacteria to remediate stress by modulating gene expression and is necessary for *C. rodentium* pathogenesis in mice. To investigate genes in the Cpx regulon that may contribute to *C. rodentium* pathogenesis, RNA-Seq, SILAC, and microarray data from previous research was mined and the genes *yebE, ygiB, bssR,* and *htpX* were confirmed to be strongly upregulated by the presence of CpxRA using *lux* reporter constructs. To determine the function of these genes *in vivo,* knockout mutants were tested in C57Bl/6J and C3H/HeJ mice. Although none of the mutants exhibited marked virulence phenotypes, the Δ*cpxRA* mutant had reduced colonization and attenuated virulence, as previously determined. We also found that the absence of the Cpx ESR resulted in higher expression of the LEE master regulator, *ler.* In addition, we determined that the Δ*cpxRA* mutant had a growth defect in medium simulating the colon, as did several of the mutants bearing deletions in Cpx-upregulated genes. Overall, these results indicate that the Δ*cpxRA* virulence defect is not due to any single Cpx regulon gene examined. Instead, attenuation may be the result of defective growth in the colonic environment resulting from the collective impact of multiple Cpx-regulated genes.

## Introduction

Attaching and effacing (A/E) pathogens like enteropathogenic and enterohemorrhagic *Escherichia coli* (EPEC and EHEC, respectively) are intestinal pathogens capable of causing severe diarrhea in children as well as adults (EHEC) (1). Due to the inability of EPEC and EHEC to replicate their human pathology in mice, *Citrobacter rodentium*, a murine A/E pathogen with genetic similarities to both EPEC and EHEC is used for *in vivo* models of infection (2, 3). *C. rodentium* causes colonic crypt hyperplasia in mice which results in severe inflammation in the colon (2, 4–6). Infections are lethal for some mice like strain C3H/HeJ, while C57Bl/6 mice experience self-limiting infections (7, 8). EPEC, EHEC, and *C. rodentium* all harbor the locus of enterocyte effacement (LEE) pathogenicity island that contains five operons (LEE1-5) encoding key proteins like intimin, Tir, and a type III secretion system (T3SS) which are responsible for the hallmark lesions formed upon intimate attachment to intestinal epithelial cells (1). The LEE is regulated by numerous factors including the master transcriptional activator Ler, encoded in the LEE1 operon, which activates the expression of LEE1-5. Ler also activates expression of the transcriptional activator, GrlA, and repressor, GrlR, encoded in the Ler-regulated *grlRA* operon that act upon LEE1 (5). In addition to LEE-encoded regulators, there are numerous other regulators of the LEE studied in EPEC and EHEC which activate or inhibit its expression based on environmental stressors sensed by the cell (9, 10).

A stress response that has been shown to be required for *C. rodentium* colonization and virulence is the Cpx envelope stress response (ESR) (11–13). The Cpx ESR is regulated by a two-component system (TCS) which consists of the sensor histidine kinase CpxA and the response regulator CpxR (14–16). In addition, it’s activity can be altered with two auxiliary signaling proteins, the inhibiting periplasmic protein CpxP and the activating outer membrane lipoprotein NlpE (16–18). While the sensing mechanism used by CpxA is poorly defined, inducing cues of the Cpx ESR include alkaline pH, salt, antimicrobial peptides, and misfolded inner membrane and periplasmic proteins (14, 16, 19–23). Upon activation, CpxA autophosphorylates and transfers a phosphoryl group to CpxR which then binds to a consensus sequence upstream of Cpx regulon members leading to altered expression. The CpxR binding consensus sequence 5’-GTAAA(N)_4-8_GTAAA-3’ is conserved to varying degrees however nucleotide deviations from the consensus sequence do not predict the relative expression of the downstream gene (20, 21, 24).

Previous studies investigating the role of the Cpx ESR in other pathogens have suggested mechanisms by which the Cpx ESR may impact, both negatively and positively, colonization and virulence. Some of these mechanisms include the negative regulation of *perC* resulting in reduced *ler* expression in EPEC, efficient expression of the bundle-forming pilus involved in initial host cell attachment in EPEC, and induction of genes required for maintaining envelope protein integrity and virulence regulation such as DegP, PpiA, and DsbA (9, 16, 25, 26). Overall, it has been concluded that the Cpx ESR facilitates adaptation to envelope stressors because it downregulates virulence genes and large protein complexes while upregulating envelope and protein modifying factors, although the specific reasons for its impact on pathogenesis have not been definitively demonstrated in most cases (16).

In *C. rodentium,* it has been concluded that the Cpx ESR is activated *in vivo* based on the observation of increased expression levels of *cpxP* (11). Given that *C. rodentium* is an A/E pathogen relying on the LEE and the encoded T3SS for virulence, it is important to note that the absence of CpxRA does not impact the secretion of T3SS effector proteins, indicating its presence has a limited impact on the overall activity of the LEE (11, 13). Differing results have been observed in terms of cell health and the presence of CpxRA when grown in virulence-inducing conditions. Thomassin *et al.* (2015) measured an extended lag phase for cells lacking CpxRA while Vogt *et al.* (2019) found no growth defect in similar conditions. The Cpx regulon in *C. rodentium* has been previously defined by microarray, RNA-seq, and SILAC proteomic analysis (12, 13). The data produced from these two studies was extensive and the impacts of numerous genes of interest on virulence have not been investigated. Giannakopoulou *et al.* (2018) determined that the auxiliary proteins, CpxP and NlpE, were not required for colonization or virulence in *C. rodentium*. On the other hand, Vogt *et al.* (2019) showed that the Cpx regulon members *degP* and *dsbA* were required for *C. rodentium* virulence in C3H/HeJ mice, however the Cpx regulation of these genes was not responsible for the virulence defect seen when *cpxRA* was absent. The Cpx regulon is extensive with the presence of CpxRA contributing to the differential expression of over 330 transcripts in *C. rodentium* (12, 13). The roles of some of the more strongly upregulated genes in both studies, such as *yebE, ygiB, bssR* and *htpX* which are investigated in this study, are not defined in *C. rodentium*.

In this study, we aimed to study the Cpx regulation of *C. rodentium* genes found in previous transcriptomic data sets as well as deduce whether they are involved in the avirulent phenotype associated with the Δ*cpxRA* mutant. We confirmed that the genes *yebE*, *ygiB, bssR* and *htpX* are upregulated in the presence of CpxRA, however, they are not required for virulence in either C57Bl/6J or C3H/HeJ mouse models. Despite this, regulation patterns influenced by the Cpx ESR as well as novel phenotypes associated with select mutants were uncovered. Using *lux* reporter genes, we confirmed reduced expression of *ler* over time in the presence of the Cpx ESR in buffered DMEM of varying glucose concentrations. Finally, we showed the activation of the Cpx ESR by high-glucose DMEM and determined that simulated colonic fluid (SCF) also induced the Cpx ESR as well as highlighted growth defects of the Δ*cpxRA* mutant and some of the Cpx-regulated gene mutants tested when combined with additional stressors (27). Cumulatively, our data suggest the Cpx ESR is required for proliferation in the gut environment by ameliorating stressors and ensuring the appropriate regulation of virulence genes which contributes to the colonization and pathogenesis of *C. rodentium*.

## Materials and Methods

### Bacterial Strains and Growth Conditions

Bacterial strains used are listed in Table S1. Unless otherwise indicated, cells were grown in either lysogeny broth (LB; 10 g/L tryptone, 5 g/L yeast extract, and 5 g/L NaCl), high-glucose Dulbecco’s Modified Eagles Medium with no phenol (Gibco^TM^, cat. no. 31053028) (HG-DMEM), or simulated colonic fluid first developed by Beumer et al. (1992), using ox bile in lieu of porcine bile (SCF; 6.25 g/L proteose-peptone, 2.6 g/L glucose, 0.88 g/L NaCl, 0.43 g/L KH_2_PO_4_, 1.7 g/L NaHCO_3_, 2.7 g/L KHCO_3_, and 4.0 g/L ox bile). Cultures were grown at 37°C with aeration at 225 rpm and LB agar plates were incubated at 37°C for 16-18 hours. For luminescence assays and growth curves, both HG-DMEM and SCF were buffered with 0.1M MOPS to maintain physiological pH of 7.4 and 7.0, respectively, unless otherwise indicated. SCF was prepared fresh for each experiment and filter sterilized. When required, media was supplemented with; 30 ug/ml kanamycin, 100 ug/ml ampicillin, 25 ug/ml chloramphenicol, 0.3mM diaminopimelic acid, 5% sucrose (filter-sterilized).

### Luminescence Reporter Construction

Luminescence reporters were constructed using the pNLP10 *lux-*reporter plasmid. Primers listed in Table S2 were designed to amplify ∼500bp upstream and 50 bp downstream of the translation start site apart from *ygiB*, where the amplified promoter was a ∼300 bp region. The restriction enzymes EcoRI and BamHI were incorporated into the forward and reverse primers, respectively (unless otherwise indicated). Promoter regions were amplified, cloned into pNPL10 and transformed into OneShot TOP10 chemically competent cells (Invitrogen, USA). TOP10 colonies harboring the pNLP10 plasmid were confirmed for insert presence using colony PCR with primers flanking the pNLP10 multiple cloning site as well as with Sanger sequencing. Plasmids with the correct insert were mini-prepped and transformed into electrocompetent *C. rodentium* DBS100 wild-type or mutant cells.

### Luminescence Assays

For spin down induction assays, cells were grown overnight in LB containing kanamycin in biological triplicate, then subcultured 1:100 and grown to an OD of 0.4-0.6. After approximately 2 hours growth, 1 ml of culture was centrifuged, the supernatant was removed, and the cells were resuspended in 1 ml pre-warmed induction medium (LB or HG-DMEM) containing kanamycin (T = 0). 200 ul of induced cells were transferred into a black walled clear bottom 96-well plate and incubated at 37°C shaking. For growth curve luminescence assays, cells were grown overnight in LB containing kanamycin in biological triplicate, then subcultured 1:100 directly into a black walled clear bottom 96-well plate containing LB with kanamycin and incubated at 37°C shaking. For all assays, empty wells were left between strains to prevent contaminating luminescence from adjacent wells. OD_600_ and luminescence measurements were taken after post-resuspension over time using the Victor X3 2030 multilabel plate reader (Perkin Elmer). *lux* activity was measured in counts per second (CPS) and normalized using the measured OD_600_ of the same well to accommodate for differences in cell number between cultures. Luminescence assays were repeated at least twice in biological triplicate.

Solid agar plate assays consisted of standardizing overnight cultures to an OD_600_ of 1.0 and serial diluting them to 10^-6^ in 1X Phosphate Buffered Saline (PBS). Each dilution was spotted onto LB supplemented with kanamycin and grown for 18 hours at 37°C. Luminescence was measured using a ChemiDoc MP imaging system (Bio-Rad). Assays were repeated twice with one representative plate shown.

### Strain Construction

All deletion mutants were generated using allelic exchange in the methods described by Vogt *et al.* (2019). In summary, in-frame deletion constructs were generated using overlap extension PCR and the primers listed in Table S2. Amplicons were then digested using the restriction enzymes XbaI and SphI/PaeI, ligated into pUC18, and transformed into OneShot TOP10 chemically competent cells (Invitrogen, USA). Plasmids were mini-prepped and sent for Sanger sequencing for amplicon confirmation. Once confirmed, the deletion construct was digested out of pUC18 and ligated into the suicide vector, pRE112, where it was transformed by electroporation into MFDpir cells (28). MFDpir cells containing the deletion construct underwent conjugation with *C. rodentium* DBS100 and single-crossover colonies were selected with chloramphenicol. Single-crossover colonies were confirmed by PCR using primers designed to flank the deletion site by ∼50 bp on each side (Table S2). Loss of pRE112 was determined by plating on LB -NaCl with 5% sucrose (filter-sterilized) and grown on benchtop for 2 days. Colonies that were sucrose-resistant and chloramphenicol-sensitive were screened by PCR to confirm intended deletion.

### C57Bl/6J and C3H/HeJ Mouse Infections

All animal experiments were performed under conditions specified by the Canadian Council on Animal Care and were approved by the McGill University Animal Care Committee. C57BL/6J and C3H/HeJ mice were purchased from the Jackson Laboratory (Bar Harbor, ME, USA). Five-week-old female mice were infected by oral gavage with 0.1 ml of LB medium containing 2-3 × 10^8^ colony-forming units (CFU) of bacteria. The infectious dose was verified by plating serial dilutions of the inoculum onto MacConkey agar (Difco). For survival analysis of C3H/HeJ mice, the mice were monitored daily and were killed if they met any of the following clinical endpoints: 20% body weight loss, hunching and shaking, inactivity, or body condition score of <2 (29). To monitor bacterial colonization, fecal pellets or the terminal centimeter of the colon were homogenized in PBS, serially diluted, and plated on MacConkey agar. Plates containing between 30 and 300 colonies were counted. Spleens were removed and weighed, and splenic indexes were calculated [√(weight of spleen × 100/weight of mouse)].

### Growth Curves

Strains were grown in biological triplicate overnight and were washed and standardized to an OD_600_ of 1.0 in 1X PBS. Cells were inoculated 1:100 into media aliquoted in a clear 96- well plate. Plates were read in an Epoch2 microplate reader (Biotek, USA) set to 37°C with continuous orbital shaking at a frequency of 237 cpm (4mm) at slow speed. Blank wells were subtracted from corresponding culture wells prior to calculations. Biological triplicates were averaged, and the standard deviation was calculated and indicated by error bars. Growth curves were completed at least twice with the data from one experiment shown. Susceptibility assays were prepared in the same manner with the addition of hydrogen peroxide or copper chloride to a final concentration of 1mM.

## Results

### Identification and confirmation of Cpx regulon members

Previous research done by two independent groups collected proteomic, RNA-Seq and microarray data to identify genes that were differentially expressed in the absence of CpxRA (12, 13). Using the data collected, the genes listed in Table 1 were selected for further study based on their predicted function, expression levels, and/or lack of previous investigation. *htpX* and *yebE* were selected as they had some of the highest transcript abundance changes besides *cpxP* and *yccA,* which have both been previously investigated in *C. rodentium* (13). SILAC data for *htpX* was insignificant due to the detection of only 1 peptide while *yebE* had a *P-*value of 0.08*. ygiB, malE,* and *dps* had significantly higher transcript abundances in one or both transcriptomic studies as well as peptide counts in the presence of CpxRA as indicated by bolded values (*dps* microarray; *P-*value = 0.081) (Table 1). *tolC* was also included as it is located 198 bp upstream of *ygiB* and has previously been presumed to be expressed in an operon with *ygiBC* (30). The additional maltose transporter complex genes including *malG, malK,* and *lamB* were included due to the significant increase in sequence abundance uncovered by microarray in the presence of CpxRA (12). *bssR* and *bdm* are involved in biofilm regulation and had significantly higher transcript abundance in wild-type cells in both transcriptomic studies despite an absence of detection in the SILAC data (Table 1).

**Table 1.** Mined RNA-Seq and SILAC data from Vogt et al. (2019) and microarray data collected in Giannakopoulou et al. (2018).

| Data Source | Dataset S3- Vogt *et al.* (2019) |  | Giannakopoulou *et al.* (2018) | Protein Function<sup>3</sup> |
| --- | --- | --- | --- | --- |
| Gene Name | Fold change RNA-Seq<sup>1</sup> | Fold change SILAC<sup>1</sup> | log2 fold change $\Delta cpxRA$-WT microarray<sup>2</sup> | |
| *htpX* | **12.10** | 10.77 | **-0.93** | Membrane-localized protease |
| *yebE* | **24.52** | 5.36 | -1.92 | Inner membrane protein with transmembrane domain |
| *ygiB* | **3.54** | **2.24** | **-0.99** | Outer membrane protein |
| *tolC*<sup>4</sup> | N/A | N/A | N/A | Outer membrane channel required for several efflux systems |
| *dps* | **2.12** | **1.43** | -2.49 | DNA protection during starvation |
| *malE* | **2.55** | **1.26** | **-5.88** | Maltose/maltodextrin-binding periplasmic protein |
| *malG* | **2.12** | n.d. | **-5.48** | Maltose/maltodextrin transport system permease |
| *malK*<sup>4</sup> | N/A | N/A | **-4.27** | Maltose/maltodextrin import ATP-binding protein |
| *lamB* | N/A | N/A | **-5.91** | Maltoporin in the outer membrane |
| *bssR* | **3.72** | n.d. | **-3.36** | Biofilm regulator involved in catabolite repression and stress response |
| *bdm* | **2.81** | n.d. | **-2.54** | Biofilm-dependent modulation protein |

**Bolded** values indicate $P_{adj} < 0.05$ (RNA-Seq), $FDR < 0.05$ (SILAC), $t\text{-test} < 0.05$ (microarray).
<sup>1</sup>Calculated WT/ $\Delta cpxRA$
<sup>2</sup>Negative value indicates higher expression in wildtype DBS100
<sup>3</sup>Protein function as described by UniProtKB for *E. coli* K12
<sup>4</sup>Included due to potential operon leader of gene of interest

Using the *lux-*reporter plasmid pNLP10, eleven reporters were constructed and tested in LB broth as well as the virulence-inducing media, high-glucose Dulbecco’s Modified Eagles Medium, which lacked phenol red and was buffered with 0.1M MOPS (HG-DMEM) (21, 31, 32). While some of these genes, namely *yebE* and *htpX,* have had Cpx-dependent expression confirmed experimentally in previous studies using other bacterial strains, their expression has never been studied in *C. rodentium* DBS100 (20, 21, 33, 34). Interestingly, the re-suspension of log-phase cells in HG-DMEM activated the Cpx ESR as seen by the near 10-fold increase in *cpxP-lux* activity relative to re-suspension in LB indicating that HG-DMEM is a stressful condition for *C. rodentium* growth (Figure 1A). Of the 11 reporters tested, *yebE, ygiB, bssR,* and *htpX* were significantly upregulated in the presence of CpxRA in both LB broth and HG-DMEM (Table S3, Figure 1A). The *yebE-lux* reporter was the most dependent on the Cpx ESR for its expression as its activity was barely detectable in both LB and HG-DMEM in the absence of CpxRA relative to its presence (Figure 1A, Table S3). In addition, similar to the *cpxP-lux* reporter, the *yebE-lux* construct was not differentially expressed in HG-DMEM relative to LB in the absence of CpxRA, unlike the other reporters tested, which exhibited higher levels of expression in HG-DMEM even without CpxRA (Figure 1A, Table S3).While we observed that the expression of *ygiB-, bssR-,* and *htpX-lux* reporters was less affected by the absence of CpxRA compared to *yebE-lux*, expression was still higher in the presence of CpxRA in both LB and HG-DMEM relative to its absence (Figure 1A). Finally, each gene of interest, except for *yebE,* had increased luminescence in wild-type cells when grown on solid LB agar relative to Δ*cpxRA* cells (Figure 1B, Figure S1A). The difference between wild-type and Δ*cpxRA yebE-lux* expression on LB agar was not visible as its basal expression was likely not detectable compared to the other reporters under these conditions.

**Figure 1.**
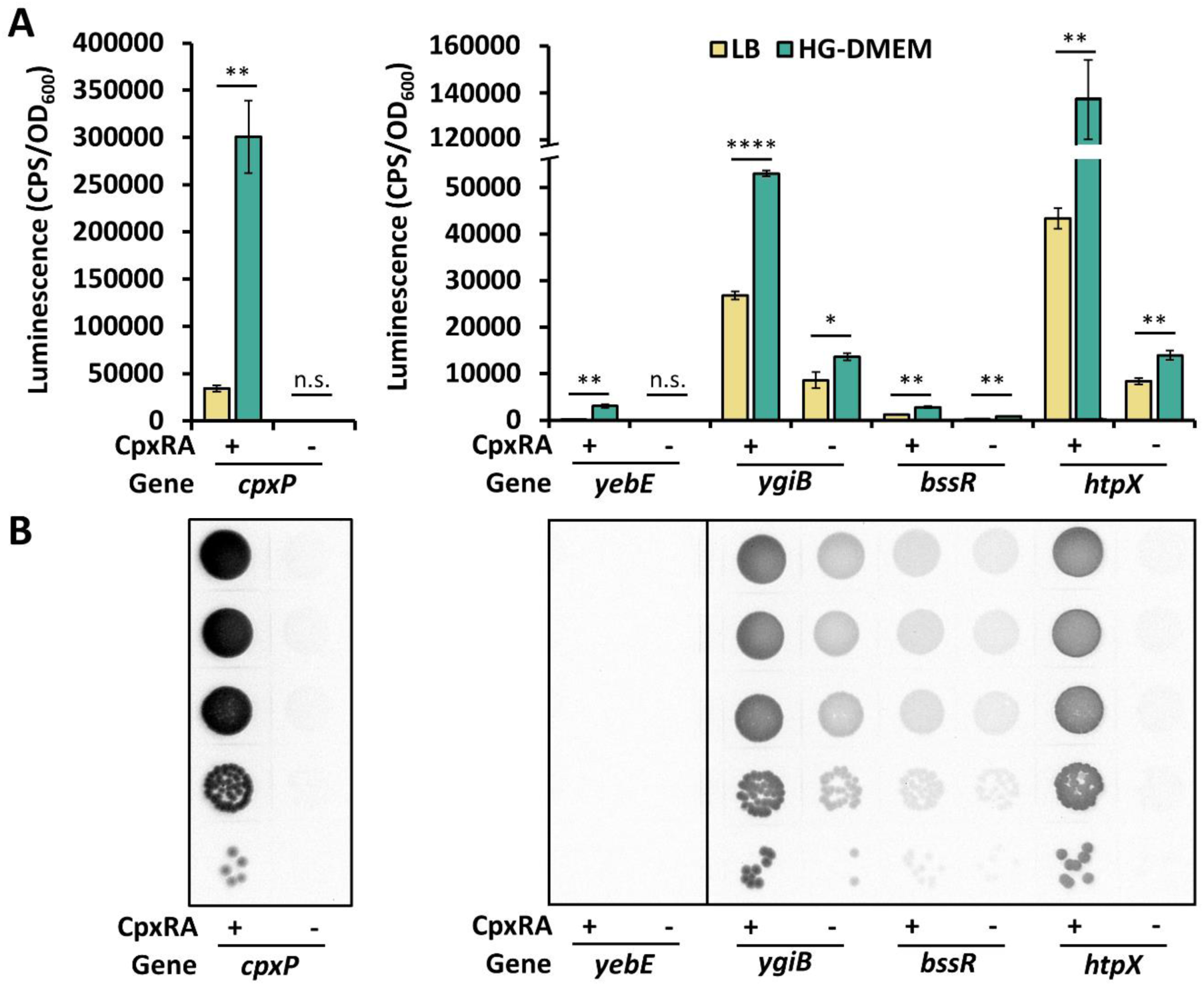
Confirmation of upregulated genes in the presence of CpxRA using *lux*-reporter assays in both LB and HG-DMEM. (A) Strains harboring *lux*-reporter plasmids were grown overnight then subcultured 1:100 in LB broth supplemented with kanamycin and grown to an OD_600_ of 0.4-0.6 at 37°C shaking. Two 1 ml aliquots of each subculture were spun down and resuspended in either LB (yellow) or HG-DMEM buffered with 0.1M MOPS (teal). The resuspended cells were added into a black walled 96-well plate and incubated at 37°C shaking. Luminescence was measured 1-hour post-resuspension in the presence or absence of CpxRA for the positive control, *cpxP*, and genes of interest *yebE*, *ygiB*, *bssR*, and *htpX* whose promoter regions were cloned upstream of the *luxCDABE* genes in the plasmid, pNLP10. Luminescence was measured in counts per second (CPS) and calculated relative to OD_600_. Data represents the mean and standard deviation of three biological replicate cultures. The asterisks indicate a statistically significant difference between the mean luminescence produced in LB compared to HG-DMEM for each strain and reporter tested (\**P* < 0.05, \*\**P* < 0.01, \*\*\*\**P* < 0.0001, n.s. not significant, Student’s *t*-test). (B) Wildtype and ΔcpxRA strains harbouring *lux*-reporter plasmids for each gene of interest were grown overnight, standardized to OD_600_ 1, serially diluted to 10^-6^, spotted on LB supplemented with kanamycin and imaged after 18 hours of growth at 37°C. Luminescence was measured using a ChemiDoc MP imaging system (Bio-Rad). All assays (A and B) were completed at least twice, with one representative experiment shown.

*tolC* was originally included in this study due to its proximity to the predicted translation initiation site of downstream *ygiB* and the presumption that it resides in an operon with *ygiBC* (30). To confirm whether these genes were in an operon together, their expression in both LB and HG-DMEM (without MOPS) was measured over 6 hours. From this, we determined there was no measurable difference in the expression of *tolC* in the presence of CpxRA in LB relative to its absence despite the activity of the Cpx response increasing over time as seen by the expression of *cpxP-lux* (Figure S2A). In HG-DMEM, the transcription of *tolC* reduces over time in the presence of CpxRA while increasing in its absence (Figure S2B). This is in contrast with the consistently higher-level expression of *ygiB-lux* in both media during the initial 4 hours of growth when CpxRA is present (Figure S2A-B). Finally, as indicated by Figure S2C, a putative CpxR binding site is located in between *tolC* and *ygiB,* 91 base pairs upstream from the predicted translation initiation site of *ygiB* (24). From this, we can conclude that *tolC* and *ygiBC* are perhaps transcribed together as suggested by Dhamdhere *et al.* (2010), however when required, the Cpx ESR has the ability to differentially express these two genes in both conditions tested.

To determine how the four selected genes of interest, *yebE, ygiB, bssR,* and *htpX,* are being expressed over time relative to the phases of cellular growth, *lux-*reporter activity was monitored for 12 hours from initial inoculation to stationary phase growth in LB. Interestingly, *yebE-lux* activity followed a near identical trend to the expression of the positive control *cpxP-lux* reporter in both wild-type and Δ*cpxRA* cells (Figure 2A). This suggests that the expression of *yebE-lux* is almost entirely dependent on the Cpx ESR or at least utilizes the same mechanisms that are responsible for the expression of *cpxP,* furthering curiosity as to its proposed function in terms of stress response and overall cell health. In addition, *ygiB-lux* and *bssR-lux* reporters showed a similar trend of expression to that of *cpxP-lux* in wild-type cells with a lower level of activity in Δ*cpxRA* cells (Figure 2B-C). It is evident for both *ygiB* and *bssR* that, unlike *yebE,* there are other regulators for these genes outside of CpxRA as they both had measurable basal expression, well above the low background levels seen for the *cpxP-* and *yebE-lux* reporters in Δ*cpxRA* cells. In addition, the peak of luminescence for *cpxP-, yebE-, ygiB-,* and *bssR-lux* in wild-type cells was reached after approximately 7-8 hours growth which coincides with late exponential phase (Figure 2A-C, Figure S3). This supports previous studies which have shown that the Cpx ESR is most active during late exponential and early stationary phase (35, 36). On the other hand, *htpX-lux* had higher luminescence in wild-type cells and was consistently expressed over the measured 12 hours of growth suggesting that its expression is less dependent on growth phase (Figure 2D). It should however be noted that the presence of CpxRA allowed for the maintenance of *htpX-lux* activity throughout late exponential and stationary phase as opposed to the steady decline in luminescence seen in the Δ*cpxRA* cells (Figure 2D, Figure S3).

**Figure 2.**
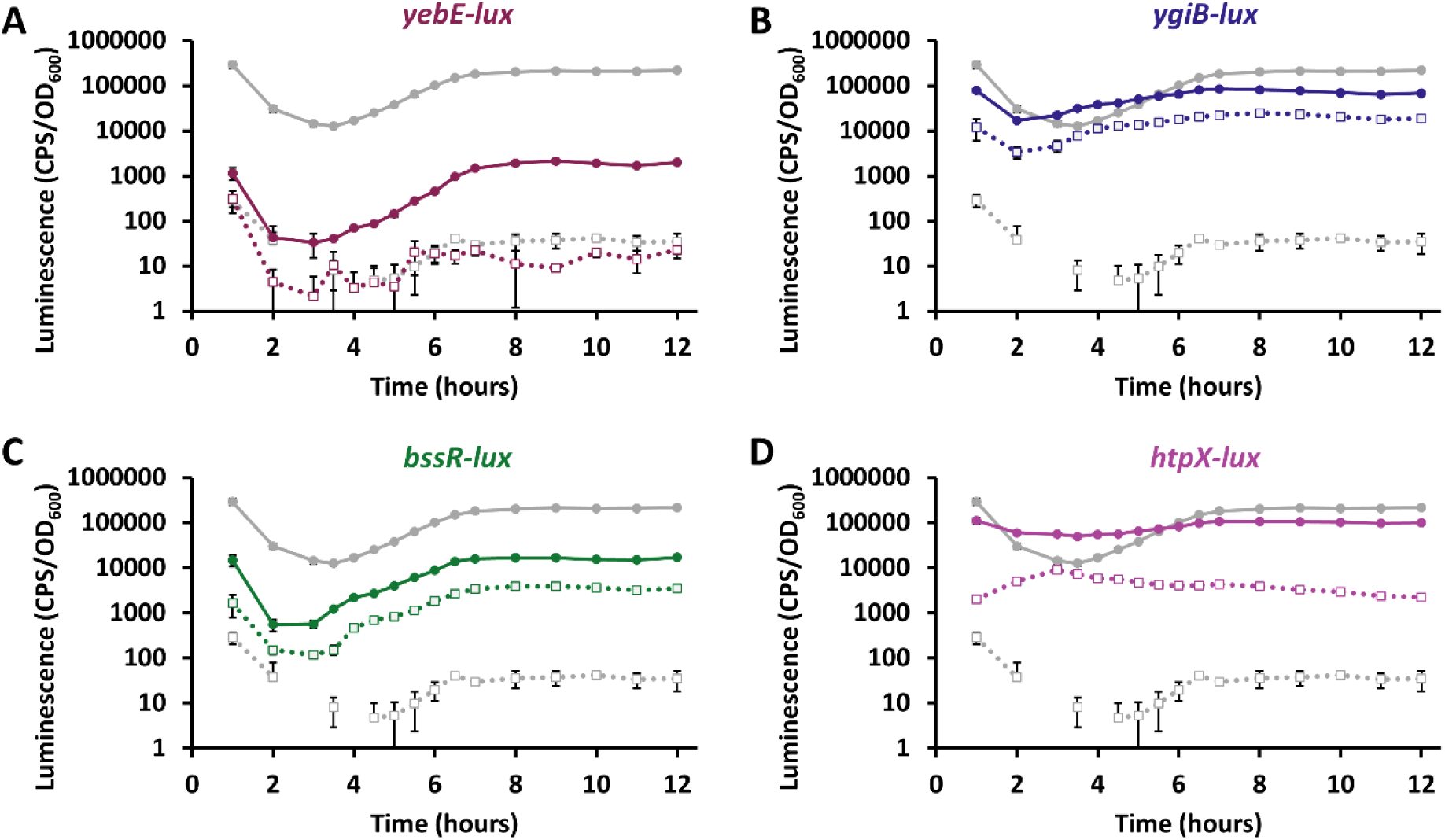
Luminescence measuring gene expression over time as compared to the activity of the Cpx ESR. Strains harboring *lux*-reporter plasmids were grown overnight and inoculated 1:100 in LB broth supplemented with kanamycin in a black walled 96-well plate and incubated at 37°C shaking. Luminescence was measured every hour for 12 hours as well as every 30 mins between 3 and 7 hours starting at 1-hour post-inoculation. The positive control *cpxP-lux* (gray) in either wildtype (line with filled circle) or *ΔcpxRA* (dashed line with empty square) backgrounds is plotted alongside the genes of interest (A) *yebE* (dark purple), (B) *ygiB* (blue), (C) *bssR* (green), and (D) *htpX* (light purple). Luminescence was measured in counts per second (CPS) and is shown relative to OD_600_. Data represents the mean and standard deviation of three biological replicate cultures. Gaps in data represent time points where no luminescence was detected thus could not be plotted on the log scale. The experiment was done twice with the results from one representative experiment shown.

### The activity of the Cpx ESR is influenced by the absence of *htpX* and growth conditions

Provided our evidence indicates *yebE, ygiB, bssR,* and *htpX* rely on the presence of CpxRA for proper expression, we questioned whether the absence of these genes would impact the envelope stress experienced by cells thus altering the activity of the Cpx ESR. Using allelic exchange, *C. rodentium* knockout mutants were generated for *yebE, ygiB, bssR,* and *htpX.* These strains were transformed with the reporter *cpxP-lux* and grown in LB or HG-DMEM where the luminescence was measured. Our results indicate that only Δ*htpX* cells had significantly increased expression of *cpxP*, supporting previously reported findings observed in *E. coli* K-12 strain MC4100 (Figure 3A-B, Figure S1B) (33). This could be seen on LB agar plates as well as in both LB broth and HG-DMEM at an increase of 2.1- and 1.9-fold, respectively (Figure 3A-B, Figure S1B).

**Figure 3.**
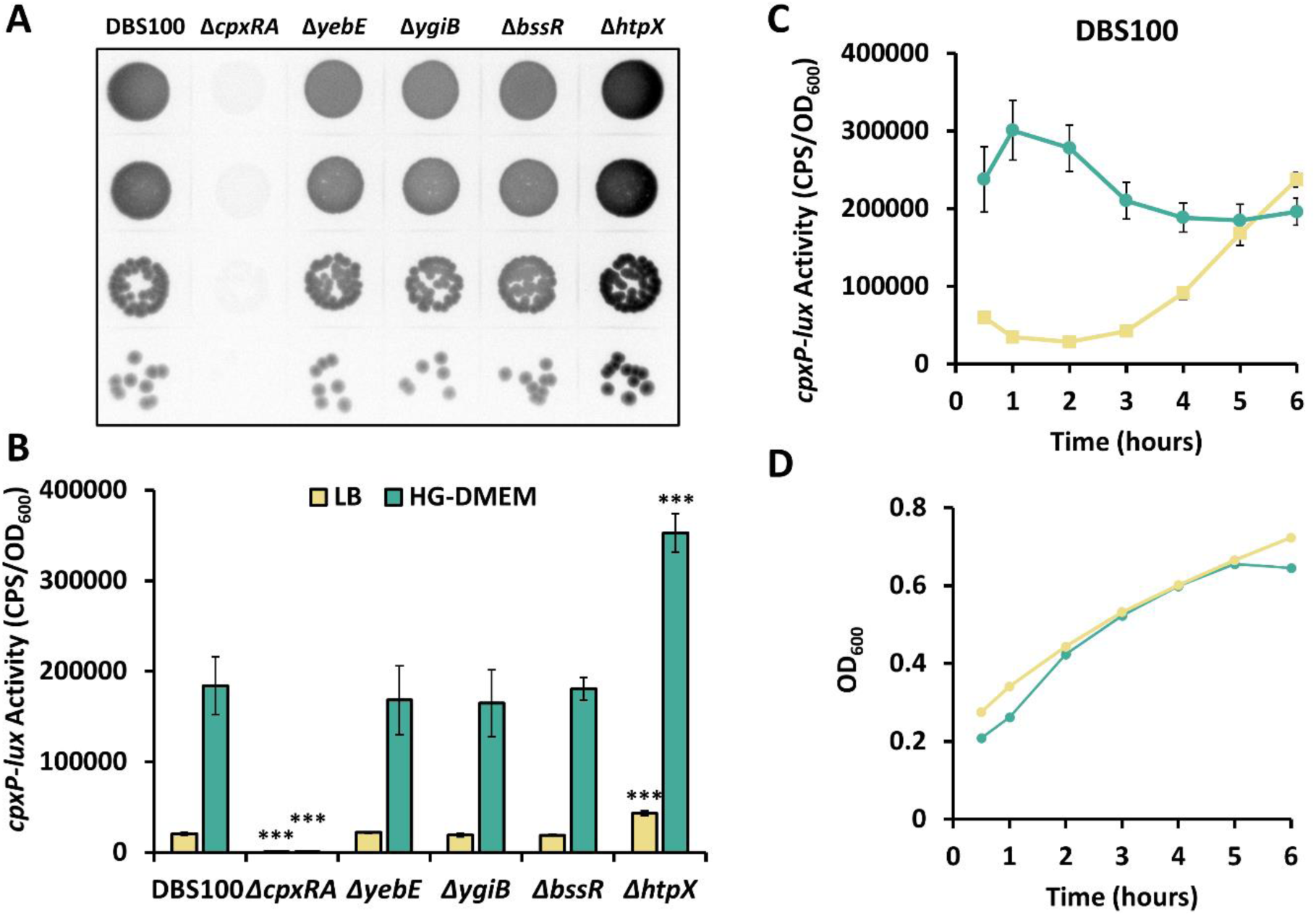
CpxRA is induced in the absence of *htpX* while the expression of *cpxP-lux* over time is increased and altered in HG-DMEM relative to LB. (A) Wildtype and mutant strains harbouring *cpxP-lux* reporter plasmids were grown overnight, standardized to OD 1, serially diluted, spotted on LB supplemented with kanamycin and imaged after 18 hours of growth at 37°C. Luminescence was measured using a ChemiDoc MP imaging system (BioRad). The experiment was repeated twice with one representative plate shown. (B-C) Strains harboring *cpxP-lux* reporter plasmids were grown overnight then subcultured 1:100 in LB broth supplemented with kanamycin and grown to an OD of 0.4-0.6 at 37°C shaking. Two 1 ml aliquots of each subculture were spun down and resuspended in either LB (yellow) or HG-DMEM buffered with 0.1M MOPS (teal). The resuspended cells were added into a black walled 96-well plate and incubated at 37°C shaking. In (B) *cpxP-lux* activity was measured 1-hour post-resuspension in wild-type, Δ*cpxRA,* Δ*yebE,* Δ*ygiB,* Δ*bssR,* and Δ*htpX C. rodentium* DBS100 strains while in (C) *cpxP-lux* activity was measured in wild-type cells every hour for 6 hours starting at 0.5 hours post-resuspension and is shown alongside (D) the respective growth curves for each media. Luminescence was measured in counts per second (CPS) and calculated relative to OD_600_. Data represents the mean and standard deviation of three biological replicate cultures. The asterisks (***) indicate a statistically significant difference from the wildtype DBS100 strain in the same media type (*P* < 0.001, one-way ANOVA with Dunnett’s multiple comparison test).

Unexpectedly, it was determined the expression pattern over time of *cpxP-lux* and by extension, the activity of the Cpx ESR, differs vastly depending on the medium used for growth (Figure 3C). When the same subculture was split and re-suspended in either LB or HG-DMEM, despite having similar growth trends over 6 hours, *cpxP-lux* activity increased over time when grown in LB but decreased over time in HG-DMEM (Figure 3C-D). In addition, while having a less substantial impact on the activity of the Cpx ESR, cultures that were grown shaking in LB induced *cpxP-lux* more so than static cultures, whereas the opposite was true for HG-DMEM cultures where static cultures had higher *cpxP-lux* activity (Figure S4A-B). These results suggest that while the Cpx ESR may be influenced by growth and is most active in late exponential or early stationary phase in LB broth, the same may not remain true for cells experiencing perhaps more stressors or other Cpx ESR inducing signals when grown in HG-DMEM.

### yebE, ygiB, bssR, and htpX are not individually required for colonization or virulence *in vivo*

With *yebE, ygiB, bssR,* and *htpX* expression confirmed to be upregulated by the presence of CpxRA, we then tested whether these genes were required for the colonization or virulence of *C. rodentium* which could provide a possible explanation for why the removal of *cpxRA* is detrimental to pathogenesis (11–13). In our first set of experiments, we used C57Bl/6J mice which experience a self-limiting form of disease to determine whether colonization was negatively impacted by any of the mutants (7). As seen in Figure 4A-C, only Δ*cpxRA* cells could not consistently colonize to the same level as wild-type and the other mutants (Day 4; *\*P*<0.05, Day 9 and 12; *\*\*P*<0.01, Mann-Whitney U Test). While the Δ*yebE* mutant exhibited a slight lag in colonization levels on day 9 (\**P*<0.05) and the Δ*bssR* mutant showed an increase in colonization of the colon on day 12 (\**P*<0.05), these minor statistical significances are not reflected in the degree of disease-state measured using a splenic index (Figure 4D). All strains caused a similar level of disease relative to wild-type except for Δ*cpxRA*, which had a significantly lower splenic index indicating attenuated virulence, thus confirming the results of previous studies (11, 12).

**Figure 4.**
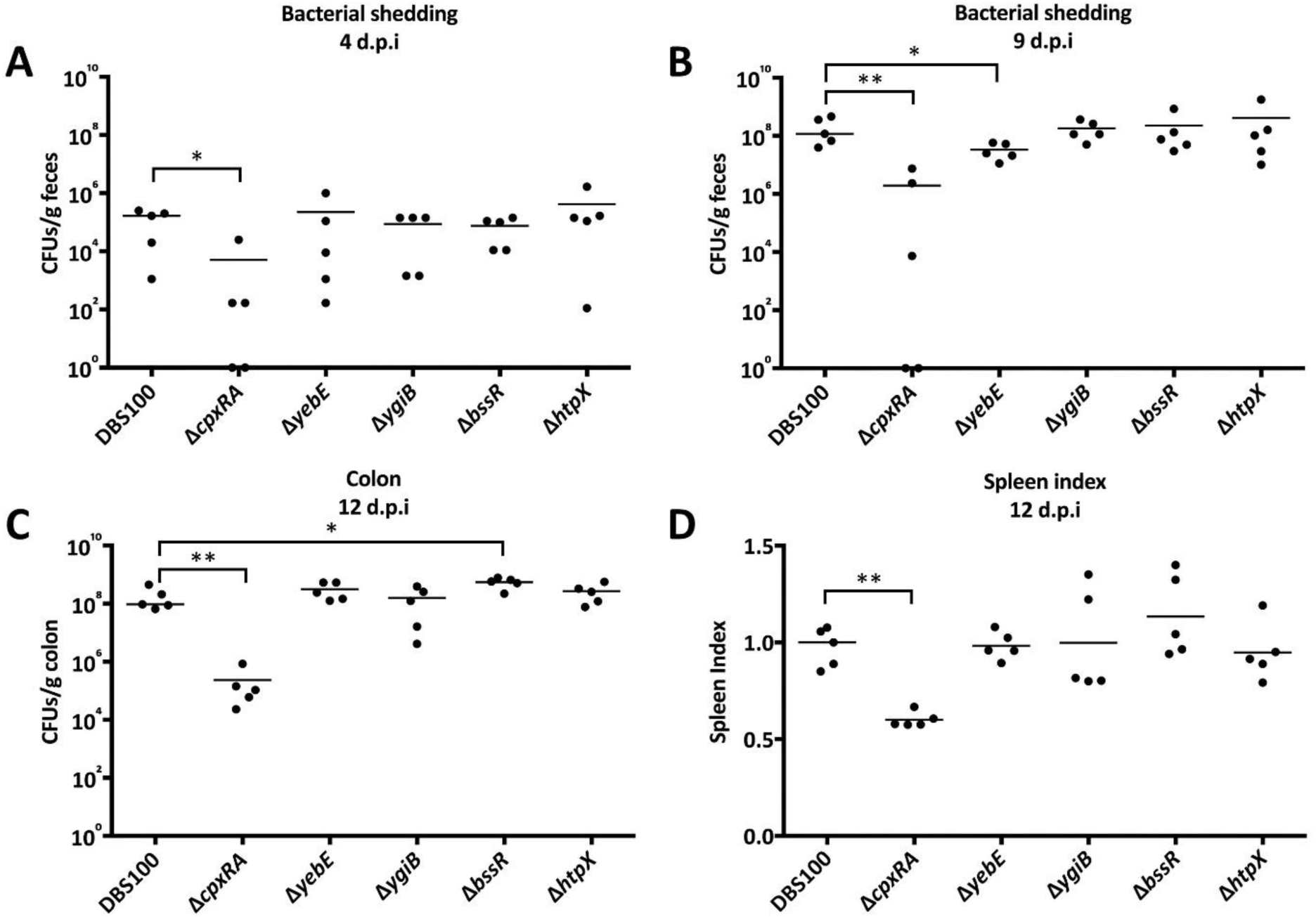
Colonization defect and reduced disease only associated with Δ*cpxRA* in C57Bl/6J mice. (A-C) Bacterial burden, per gram of feces at days 4 and 9 post-infection or the terminal centimeter of the colon on day 12, was measured in colony forming units (CFUs) by selective plating on MacConkey agar. (D) Spleens were harvested from euthanized mice and the spleen index was determined relative to mice infected with wild-type *C. rodentium.* (A-D) Horizontal lines indicate the median of n=5 mice and asterisks show significant differences between mice infected with wild-type versus those infected with a mutant strain (*\*P*<0.05, *\*\*P*<0.01, Mann-Whitney U Test).

Similarly, in a C3H/HeJ mouse trial, testing disease progression and survival, only the Δ*cpxRA* mutant exhibited a colonization defect as seen by an approximate 2-fold reduction in CFUs for three mice and undetectable levels in two on day 4 post-infection (Figure 5A). In addition, the Δ*cpxRA* mutant had significant attenuation of virulence as seen by the fact that all five infected mice survived until day 30 (Figure 5B).

**Figure 5.**
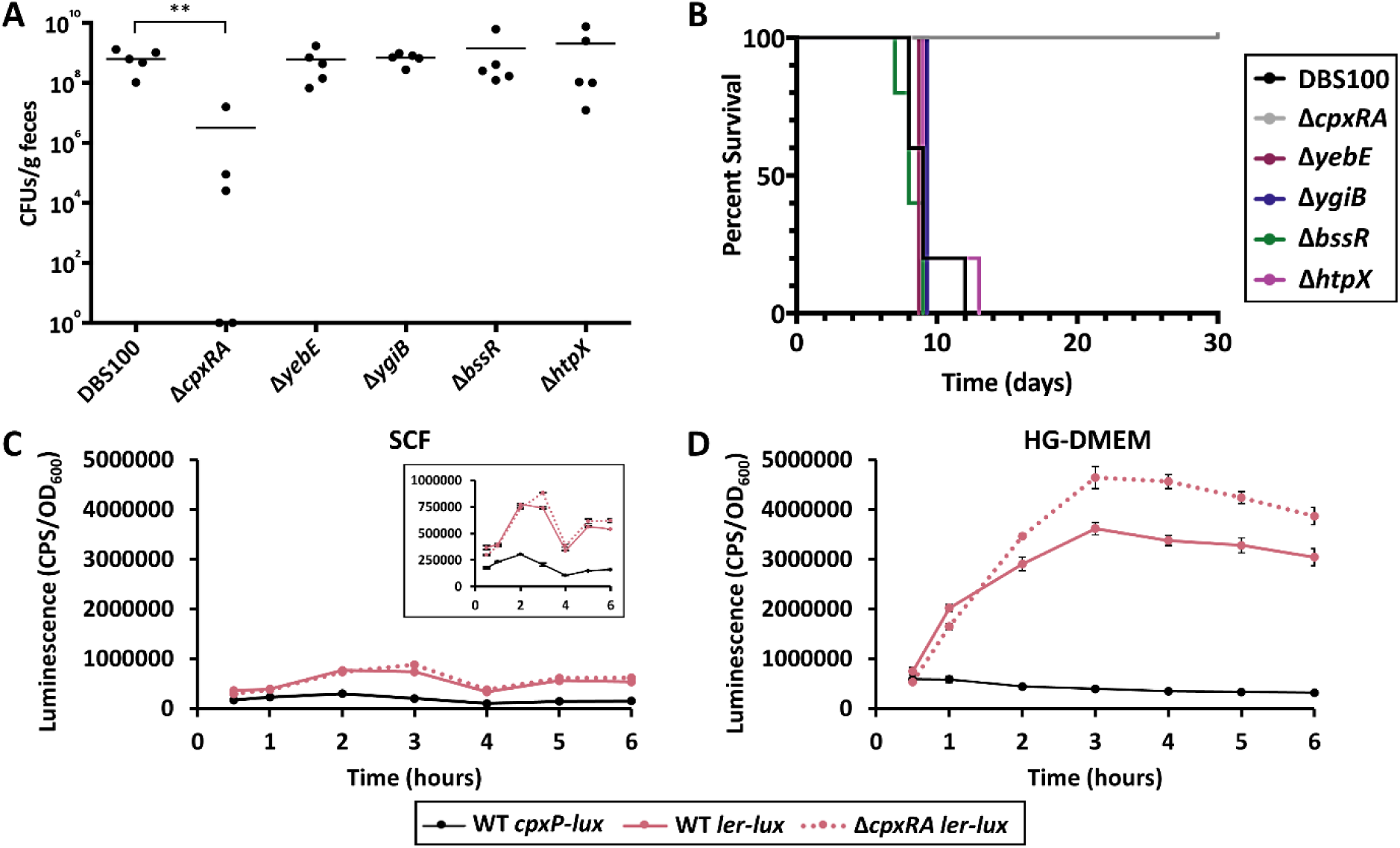
Prominent regulon members presence does not provide cause for *C. rodentium* Δ*cpxRA* attenuation in C3H/HeJ mice while Δ*cpxRA* mutants shows altered expression of LEE regulator, *ler*. (A) Colonization level was determined 4 days post-infection and measured in colony forming units (CFUs) per gram of feces. Horizontal lines indicate the median of n=5 mice and asterisks show significant differences between mice infected with wild-type versus those infected with a mutant strain (*\*\*P*<0.01, Mann-Whitney U Test). (B) Data depicts percent survival of mice infected with *C. rodentium* DBS100 strains over 30 days. Mice were euthanized if they reached any one of the following critical endpoints: 20% body weight loss, hunching and shaking, inactivity, or body condition score of <2. (C-D) Wild-type (solid line) and Δ*cpxRA* (dotted line) cells harboring *ler-lux* (pink) reporter plasmids and wild-type cells containing *cpxP-lux* (black) were grown overnight then subcultured 1:100 in LB broth supplemented with kanamycin and grown to an OD of 0.4-0.6 at 37°C shaking. Two 1 ml aliquots of each subculture were spun down and resuspended in either (C) SCF buffered with 0.1M MOPS or (D) HG-DMEM buffered with 0.1M MOPS. The resuspended cells were added into a black walled 96- well plate and incubated at 37°C statically. The plate was shaken for 30 seconds prior to measuring luminescence in counts per second (CPS) every hour for 6 hours starting at 0.5 hours post-resuspension. Reporter activity was plotted relative to OD_600,_ and data represents the mean and standard deviation of three biological replicate cultures.

### The Cpx ESR reduces the transcription of the LEE master regulator in HG-DMEM

Previous studies have demonstrated the *C. rodentium* Δ*cpxRA* mutant does not have a significantly altered T3SS secreted protein profile, while there is reduced transcription of LEE operons upon activation of CpxRA and reduced expression of translocator proteins in EPEC and EHEC (11, 13, 37, 38). Utilizing *lux* reporter genes, we asked whether the attenuation of the *C. rodentium* Δ*cpxRA* mutant could be in part due to altered transcription of the LEE operon by measuring the luminescence of a *ler-lux* reporter. When cells were grown statically in simulated colonic fluid (SCF), a medium that simulates the lumen of the colon, the expression of *ler* was independent from the presence of the Cpx ESR (Figure 5C) (27). When cells were grown statically in the virulence-inducing condition, HG-DMEM, *ler* expression was higher in Δ*cpxRA* mutant cells indicating that the presence of the Cpx ESR reduces the expression of *C. rodentium’s* primary LEE regulator (Figure 5D).

### Cpx-regulated genes impact *C. rodentium* fitness in simulated colonic fluid (SCF)

Previous research has shown differing results with regard to the effect of removing the Cpx ESR on *C. rodentium* growth. Vogt *et al.* (2019) showed no growth defects associated with a *ΔcpxRA* mutant in shaking LB broth or static high-glucose DMEM with 5% CO_2_. On the other hand, although the exact nature of the culture conditions used are unclear, Thomassin *et al.* (2015) found that *C. rodentium* Δ*cpxRA* cultures had a longer lag phase in DMEM but eventually would grow to a comparable OD to the wild-type and complemented strains. In this study, all the mutants tested grew comparably in LB, however in buffered HG-DMEM, all the mutants grew to a reduced level relative to wild-type cells, with the most significant reduction experienced by Δ*cpxRA* cells (Figure 6A-B). One predominant issue to note with *C. rodentium* in HG-DMEM is an overall poor growth phenotype as been by the OD maximum of 0.2 (Figure 6B). To circumvent this as well as to mimic the *in vivo* conditions experienced by *C. rodentium* cells during colonization, the strains were grown in simulated colonic fluid (SCF), first developed by Beumer *et al.* (1992). Interestingly, when comparing unbuffered and buffered growth in SCF, it becomes evident that while unstable pH is toxic to Δ*cpxRA* cells, resulting in zero growth, there is also a growth defect evidenced by a longer lag phase in MOPS buffered SCF at pH 7 (Figure 6C-D). Importantly, when pH is controlled in buffered SCF, only the Δ*cpxRA* cells grow significantly different from wild-type cells, mimicking the colonization defect seen in both C57Bl/6J and C3H/HeJ mice (Figure 4A-C, Figure 5A, Figure 6D). The Δ*cpxRA* cells also exhibited severe growth defects when grown in buffered SCF and challenged with the presence of either oxidative or copper stress, suggesting an extreme sensitivity to sub-inhibitory levels of stressors in the colonic environment (Figure 7). An additional observation to be noted in unbuffered SCF is that both the Δ*ygiB* and Δ*htpX* mutants experienced an extended lag phase as well as had increased variability between biological replicates (Figure 6C). This could indicate a susceptibility to unstable pH as well as suggests a contributing factor to the pH sensitivity experienced by Δ*cpxRA* cells (Figure 6C).

**Figure 6.**
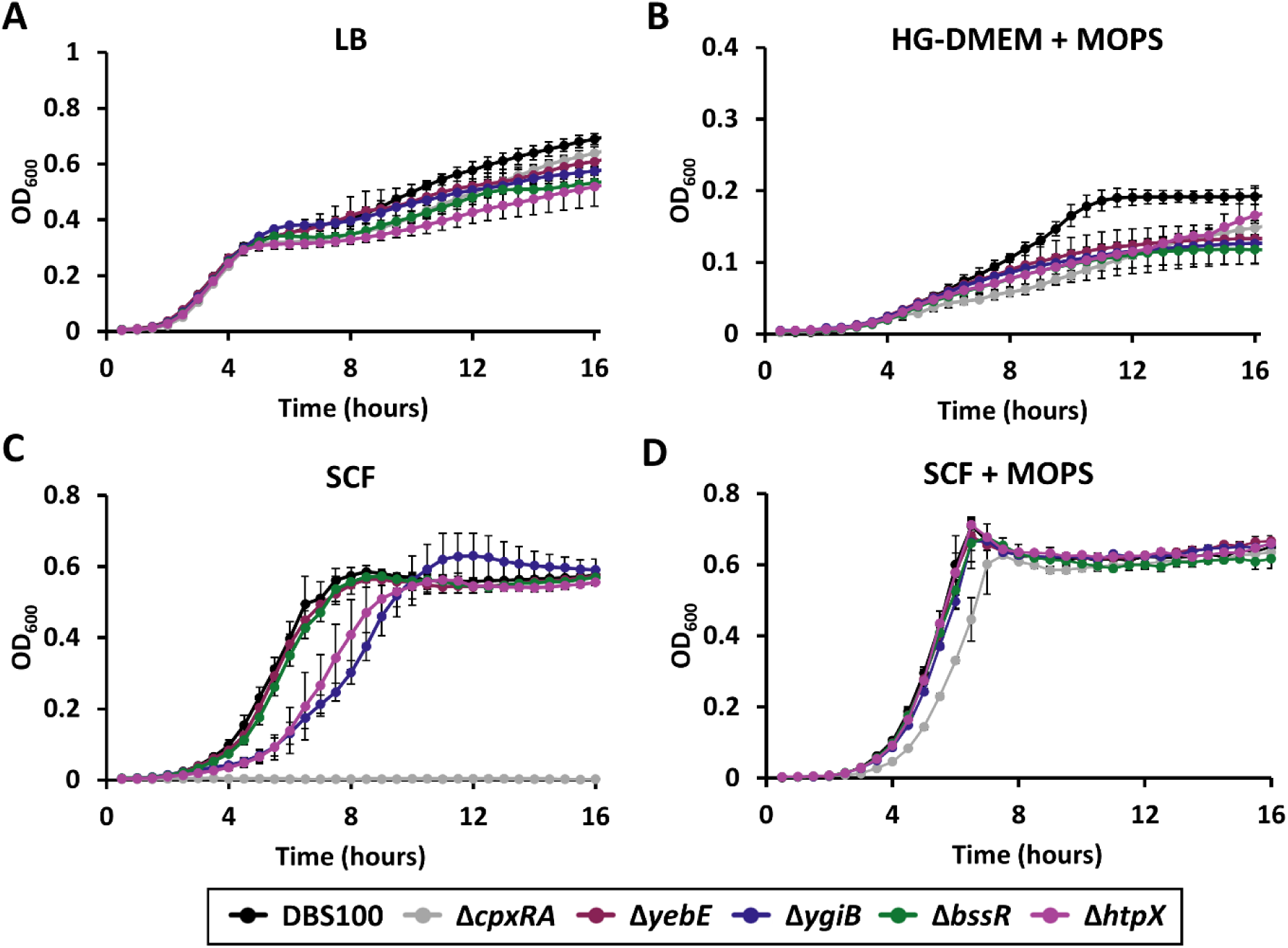
Simulated colonic fluid (SCF) promotes growth more so than HG-DMEM whilst indicating fitness defects in Δ*cpxRA*, Δ*ygiB*, and Δ*htpX*. Strains were grown overnight in LB at 37°C shaking. Cells were washed in PBS, standardized to OD_600_ 1.0, and inoculated 1:100 into (A) LB, (B) HG-DMEM with 0.1M MOPS, (C) SCF, and (D) SCF with 0.1M MOPS in a clear 96-well plate. OD_600_ measurements were taken every 30 minutes for 18 hours using an Epoch2 microplate reader (BioTek, USA). Blank well measurements were subtracted from sample OD_600_ measurements. Data represents the mean of three biological replicates and the error bars indicate the standard deviation. The experiment was completed twice with the data from one experiment shown.

**Figure 7.**
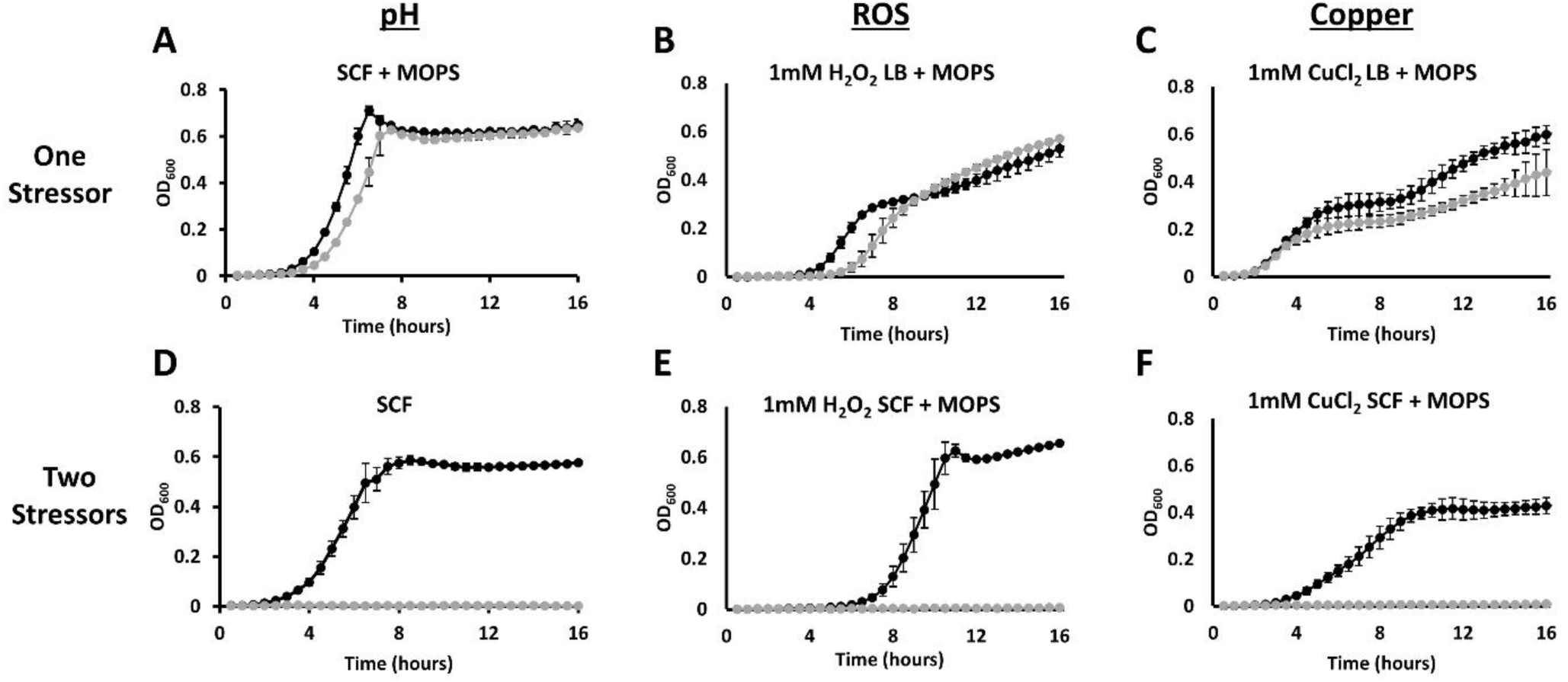
Simulated colonic fluid (SCF) highlights severe fitness defects in Δ*cpxRA* cells in response to sub-inhibitory levels of pH, oxidative, and copper stress. Strains were grown overnight in LB at 37°C shaking. Cells were washed in PBS, standardized to OD_600_ 1.0, and inoculated 1:100 into (A, D, F) SCF with 0.1M MOPS (B, C) LB with 0.1M MOPS or (D) SCF in a clear 96-well plate. Sub-inhibitory levels of the stressors (B, E) 1mM H_2_O_2_ to simulate oxidative stress and (D, F) 1mM CuCl_2_ for copper stress were added to cultures containing wild-type (black) or Δ*cpxRA* (grey) cells. OD_600_ measurements were taken every 30 minutes for 18 hours using an Epoch2 microplate reader (BioTek, USA). Blank well measurements were subtracted from sample OD_600_ measurements. Data represents the mean of three biological replicates and the error bars indicate the standard deviation. The experiments were completed three times with one representative experiment of each stressor shown.

Due to the susceptibility to stressors exhibited by the Δ*cpxRA* mutant in SCF, we investigated whether our Cpx-regulated gene mutants also experienced increased sensitivity when in the presence of oxidative stress. While growth was observed in the presence of oxidative stress in buffered LB, the knockout mutants Δ*yebE,* Δ*ygiB,* and Δ*htpX* all exhibited varying degrees of susceptibility to sub-inhibitory levels of hydrogen peroxide when grown in buffered SCF, with Δ*ygiB,* and Δ*htpX* mutants showing the greatest defects (Figure 8). Finally, a reduction in OD indicating cell lysis was also observed in SCF buffered with MOPS as cells transitioned from exponential to stationary phase (Figure 6D, Figure 8B-D). The cause of this reduction requires further study. These results indicate that SCF mimics the colonization defect of the *ΔcpxRA* mutant measured *in vivo* which suggests that it may be a better medium to use when evaluating potential *in vivo* growth phenotypes. In addition, SCF appears to be a medium that can highlight subtle growth defects that might not be as easily detected in an animal model as seen with the Δ*yebE,* Δ*ygiB,* and Δ*htpX* mutants.

**Figure 8.**
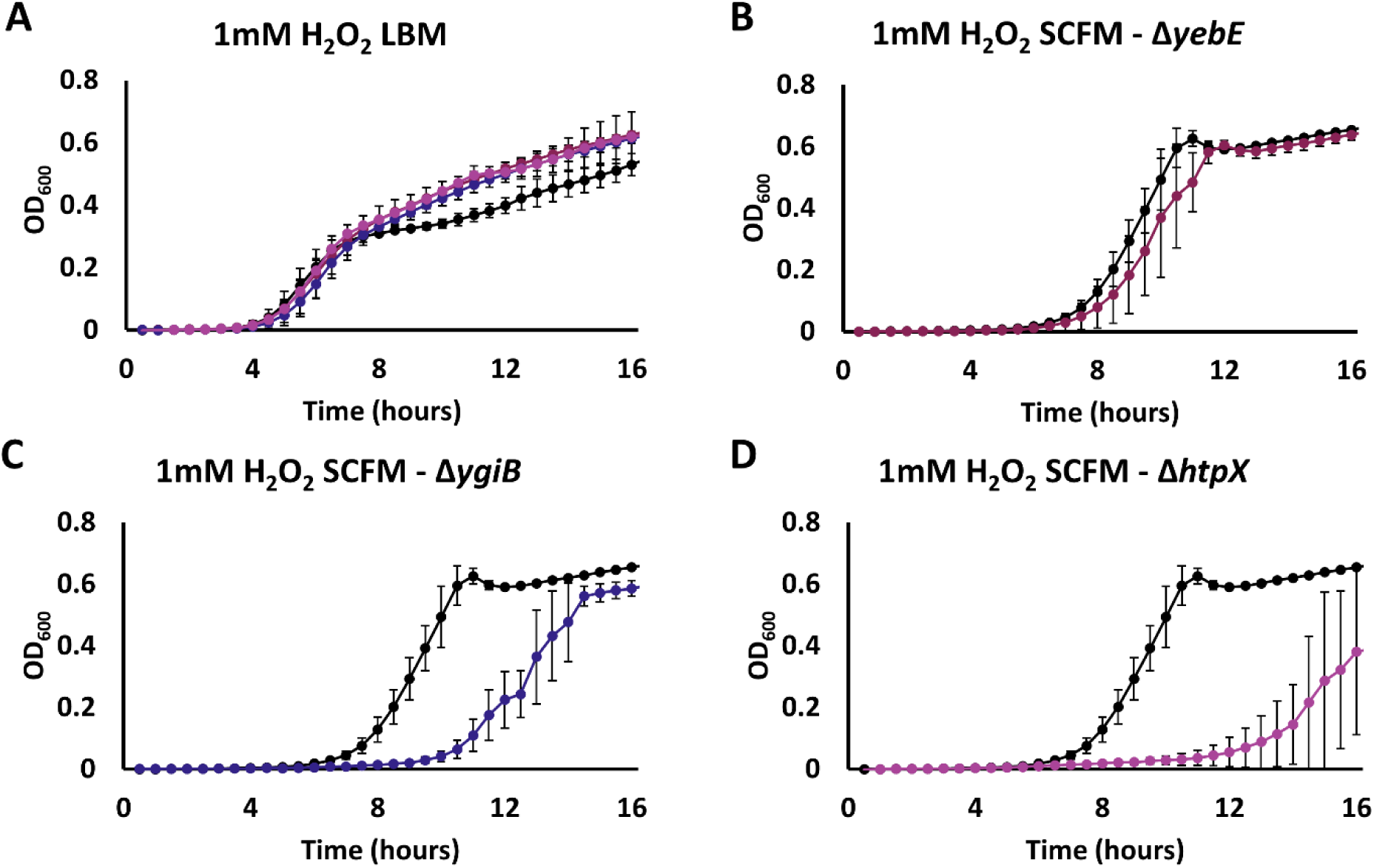
Δ*yebE,* Δ*ygiB* and Δ*htpX* cells experience varying levels of fitness defects in simulated colonic fluid (SCF) when grown in sub-inhibitory levels of oxidative stress. Strains were grown overnight in LB at 37°C shaking. Cells were washed in PBS, standardized to OD_600_ 1.0, and inoculated 1:100 into (A, D, F) SCF with 0.1M MOPS (B, C) LB with 0.1M MOPS or (D) SCF in a clear 96-well plate. Sub-inhibitory levels of the stressors (B, E) 1mM H_2_O_2_ to simulate oxidative stress and (D, F) 1mM CuCl_2_ for copper stress were added to cultures containing wild-type (black) or Δ*cpxRA* (grey) cells. OD_600_ measurements were taken every 30 minutes for 18 hours using an Epoch2 microplate reader (BioTek, USA). Blank well measurements were subtracted from sample OD_600_ measurements. Data represents the mean of three biological replicates and the error bars indicate the standard deviation. The experiments were completed three times with one representative experiment of each stressor shown.

## Discussion

### Induction of the Cpx ESR over time is condition-specific

Throughout this study, we identified numerous growth conditions which influenced the activity of the Cpx ESR. HG-DMEM is commonly used for inducing the expression of virulence factors *in vitro* (31, 32). A previous study published microarray data indicating that HG-DMEM also moderately induced *cpxP* gene expression in EPEC overexpressing NlpE though they did not comment on this result being DMEM-dependent (34). Whilst DMEM has been used in previous Cpx-related studies to induce virulence gene expression, to our knowledge this is the first study to distinctly demonstrate a strong induction of the Cpx ESR in HG-DMEM relative to LB (Figure 1A) (13, 25, 26, 34, 37, 38). The Cpx ESR is associated with monitoring proper membrane protein biogenesis, the repression of virulence factors, and maintaining cell wall integrity (16). Therefore, its increased level of activity in HG-DMEM highlights the importance of stringent regulation of envelope functions under conditions where virulence factor expression is upregulated and suggests that the proper coordination of virulence factor production with envelope homeostasis may be just as important as their presence for pathogenesis *in vivo*.

Interestingly, while previous studies have shown that Cpx ESR activity is highest in late exponential or early stationary phase in LB, we found the activity of the Cpx ESR in HG-DMEM, indicated by the expression of the *cpxP-lux* reporter, was highest during log phase growth and reduced over time (Figure 3C-D) (35, 36). If HG-DMEM was a simple activator of the Cpx ESR, one would expect the pattern of expression for the positive control, *cpxP,* to remain the same albeit with the luminescence levels increased, however this was not observed (Figure 3C). These observations over time indicate the dynamic nature of the Cpx ESR and how the integration of numerous, likely intrinsic and extrinsic, signals can alter its activity. While single time points can indicate activation or repression by the Cpx ESR, collecting data over time allows for a further understanding of the reliance a gene’s expression has on the Cpx ESR throughout growth as evidenced by Figure 2A-D.

In addition, not only did the media affect the activity levels of the Cpx ESR over time but also cultures grown in shaking versus static conditions impacted the level of Cpx activity in a media-dependent manner (Figure S4). Expression of the *cpxP-lux* reporter was influenced differently by a shaking versus static culture depending on the media used as opposed to the speed of growth, as both cultures grew faster in shaking conditions (data not shown). Previous studies have shown that static and shaking cultures can impact gene regulation and resulting phenotypes. In *Salmonella enterica*, it has been shown that the expression of *hilA* is CpxA- dependent in low pH and it’s expression is increased when cultures are grown statically (39). In uropathogenic *E. coli,* the agglutination titer was opposite for wild-type and strains lacking the small RNA, RyfA, when either grown shaking or statically in both LB and human urine (40). Therefore, our study highlights the importance of culture conditions on the activity of the Cpx ESR and prompts questions as to the nature of the envelope stressors in the medias tested and by extension how those conditions could impact the physiology of growing cells.

### Elucidating the regulation of uncharacterized genes by the Cpx ESR

One of the objectives of this study was to develop an understanding of the Cpx-dependent regulation for the relatively uncharacterized genes *yebE* and *ygiB.* In *E. coli,* YebE is an uncharacterized predicted inner membrane protein that shares homology with TerB-like proteins and contains putative metal binding sites (UniProtKB Accession no. P33218, GenBank CAD6012101.1). According to the protein prediction software InterProScan and TMHMM Server v. 2.0, YebE has two transmembrane domains, a linker in the cytoplasm, and a large globular protein structure in the periplasm (41–44). YebE is a widely conserved protein largely in the classes alpha-, delta- and gammaproteobacteria with homologs identified in Pseudomonales, Enterobacterales, Rhizobiales, and Burkholderiales among others, as identified by the database eggNOG 5.0 (45). Previous genome screens and microarrays have correlated the induction of *yebE* expression in *E. coli* with fluoroquinolone resistance, and in response to alkaline pH, copper stress, and UV irradiation though minimal work has been done to characterize the function of the encoded protein (20, 46, 47). *yebE* is unique in this study as its expression appears to rely largely on the presence of the Cpx ESR in the conditions tested (Figure 1A, Figure 2A). While other genes, like *ygiB, bssR,* and *htpX,* maintained expression in the Δ*cpxRA* mutant, the expression of *yebE* was mostly abolished in LB, and in wild-type *C. rodentium* closely resembled the expression pattern of *cpxP* over time (Figure 1A, Figure 2A). While we have found that YebE is not required for colonization nor virulence *in vivo*, our results indicate the encoded protein could perhaps aid in maintaining cell membrane integrity and/or mediating oxidative stress during growth in simulated colonic fluid providing direction for future study of this gene (Figure 8B).

YgiB is a predicted outer membrane lipoprotein which was originally suggested to be encoded in an operon with *tolC* (30). Despite this, our results indicate that *ygiB* is under the control of its own promoter and can be differentially expressed from *tolC* by the Cpx ESR in that *ygiB* expression is induced upon stress while *tolC* is reduced (Figure S2). Currently the only major phenotypes associated with *ygiB* include an exacerbation of Δ*tolC* growth defects in minimal media with glucose when cells are lacking the YgiBC and YjfMC proteins as well as an induction of expression during mature biofilm formation (30, 48). When grown in simulated colonic fluid in the presence of hydrogen peroxide to induce oxidative stress, Δ*ygiB* cells were unable to grow until approximately 10 hours post-inoculation where apparent suppressor mutations developed as indicated by the large deviations in growth between biological triplicates (Figure 8C). In addition, albeit slight, we found that Δ*ygiB* cells have a reduced growth rate relative to wild-type cells in the presence of copper stress when grown in SCF (data not shown). Numerous studies have shown that *tolC* is required for resistance to bile as a part of the AcrAB-TolC efflux pump in *E. coli* as well as colonization *in vivo* for various pathogenic bacteria (49–53). Given the exacerbation of the Δ*tolC* mutant growth defect with the absence of *ygiBC* demonstrated by Dhamdhere *et al*. (2010), the differential regulation of *tolC* and *ygiB* by the Cpx ESR seen in this study, and the susceptibility of Δ*ygiB* cells to additional stressors when grown in SCF containing bile, it is intriguing to suggest that perhaps YgiB complements the function of TolC in stressful conditions under the control of the Cpx ESR. In other words, the Cpx ESR may act to minimize large membrane proteins like TolC from integrating into a stressed membrane and instead upregulate YgiB to carry out similar functions in the meantime. Since evidence for the function of *ygiB* is largely lacking in current literature, these results could provide an interesting avenue for future work.

### The presence of the Cpx ESR negatively impacts the expression of master LEE regulator, *ler*

Previous studies have shown that the Cpx ESR negatively impacts the expression of virulence factors. In EPEC, overexpression of the response regulator CpxR reduced the activation of LEE1, LEE4, and LEE5 while the removal of CpxR resulted in increased expression of all five LEE operons when in a non-pathogenic *E. coli* background that lacked *ler* indicating the regulation of LEE by the Cpx was likely *ler-*independent (38). In EHEC, it has also been shown that the Cpx ESR negatively regulates virulence factors, including those that are LEE-encoded through Sigma factor 32 and the Lon protease (37). On the other hand, a recent study by Kumar *et al.* (2020), proposed a model for EHEC suggesting that CpxR upregulates the expression of *ler* directly and that serotonin is an inhibitor of the Cpx ESR which results in reduced transcription of LEE. The authors used qRT-PCR and growth in low-glucose DMEM to show that *ler* was reduced in the absence of CpxR in EHEC and was reduced in *C. rodentium* Δ*cpxA*. Due to these findings, we wished to verify the impact of the Cpx ESR on the expression of *ler* to determine whether the avirulence associated with the Δ*cpxRA* mutant could be due to reduced expression of LEE in *C. rodentium*. Unlike Kumar *et al.* (2020), our data indicates that in SCF, the impact of the Cpx ESR on *ler* expression is minimal with a slight increase in expression in the absence of the Cpx ESR (Figure 5C). We also found that in both high- or low-glucose DMEM, in static and shaking conditions, the expression of *ler* is consistently higher in the absence of the Cpx ESR (Figure 5D, Figure S5A-D). Therefore, our data suggests the colonization and virulence phenotypes observed for the Δ*cpxRA* mutant is not due to reduced expression of *ler* but perhaps could be from the overexpression of *ler* which could contribute to reduced fitness and inappropriately timed virulence mechanisms *in vivo*.

### SCF is beneficial for determining colonization efficacy and susceptibilities of mutants to stressors

While HG-DMEM has been a frequent media used to induce virulence gene expression and mimic an *in vivo* environment prior to conducting animal model experiments, we chose to investigate the fitness of our mutants in a medium relevant to the conditions present in the colon by using SCF (31, 32). SCF is a relatively uncommon media in individual pathogen fitness studies as in the past it has primarily been used for determining drug solubility and delivery systems which involve looking at the microbiota’s affect on drug release (55, 56). A more frequently used gastrointestinal fluid is simulated gastric fluid (SGF) which has been used to demonstrate acid-tolerance of pathogens like EPEC, EHEC, *Vibrio cholerae, Listeria monocytogenes,* and *Salmonella* (57–60). One study in *Salmonella* demonstrated differences in cell viability upon exposure to gastrointestinal fluids where they found gastric juice with a pH of 4 or 5 in conjunction with bile salts from simulated intestinal juice reduced cell viability greater than acidic pH alone (60). In this study we show that simulating the conditions *C. rodentium* cells face during colonization of the colon using SCF was able to uncover growth defects and susceptibilities in Δ*cpxRA* cells and the mutants of our genes of interest that would have been unidentified in LB or HG-DMEM. Of note, wild-type *C. rodentium* grew to a significantly higher OD in SCF relative to HG-DMEM and had an increased growth rate relative to that in LB suggesting that this media simulates an environment this gastrointestinal pathogen has adapted to (Figure 6B-C). In addition, the growth phenotypes in buffered SCF paralleled the *in vivo* colon colonization of each mutant thus highlighting the versatility and replicability of using simulated physiological conditions (Figure 5C, Figure 6D). Furthermore, mutant cells grown in SCF were more susceptible to extraneous stressors than wild-type *C. rodentium* which could allow for greater insight into the function of genes like *yebE* and *ygiB* which previously had no known associated growth phenotypes (Figure 8).

Interestingly, the Δ*cpxRA* mutant had an increased lag phase when grown in the presence of oxidative stress in LB while growth was abolished in SCF with H_2_O_2_ (Figure 7B and E). As detailed thoroughly in recent reviews, it is understood that *C. rodentium* utilizes aerobic respiration to outcompete host microbiota during colonization (6, 61, 62). A previous study found that deletion of the *cydAB* genes in *C. rodentium*, which are required for aerobic respiration, resulted in a severe reduction of growth *in vivo* (63). Following this, it was determined that disruption to the mitochondrial respiration of intestinal epithelial cells is largely responsible for *C. rodentium* infection causing oxygenation of the mucosal surface as opposed to solely colonic crypt hyperplasia (64, 65). The Cpx ESR has been implicated in the regulation of aerobic respiration in EPEC where it has been shown that removal of *cpxRA* reduced the oxygen consumption capabilities of cells which was attributed to problems with cytochrome *bo_3_* oxidase biogenesis or function (66). Similar conclusions have been made in *Salmonella* Typhimurium which also utilizes aerobic respiration to expand in the gastrointestinal tract and experiences colonization defects in the absence of CpxRA (67, 68).

A follow up study to Lopez *et al.* (2016) found that prior to expansion by aerobic respiration, *C. rodentium* utilizes host derived H_2_O_2_ as an electron acceptor during anaerobic respiration via cytochrome *c* peroxidase (*ccp*) (69). Both wild-type and Δ*ccp* mutant cells could survive in LB in the presence of mM concentrations of H_2_O_2_, though in the absence of *ccp* there was increased expression of the catalase-peroxidase, *katG,* suggesting that cells were experiencing higher levels of oxidative stress. In the RNA-seq data from Vogt *et al.* (2019), *katG* expression was also significantly induced in the *cpxRA* mutant grown in HG-DMEM which suggests the cells were experiencing elevated levels of oxidative stress (data not shown). Given that rapid expansion by aerobic respiration is a proposed mechanism of *C. rodentium* pathogenesis in overcoming colonization resistance and the production of reactive oxygen species by intestinal epithelial cells is an important defense mechanism, our results utilizing SCF support the implication that the Cpx ESR is required to successfully mediate encountered oxidative stressors during colonization in the gastrointestinal tract (63, 70–72).

The formulation of SCF used in this study, with the substitution of ox bile for porcine bile, was developed by Beumer *et al.* (1992) with the intentions of isolating non-culturable but viable *Campylobacter jejuni* coccoid cells. The recipe for this media proved unique from other versions of colonic fluid that have been used to study EHEC as it contained proteose-peptone, used porcine bile as opposed to bile salts, as well as lacked Bacto tryptone (73, 74). Despite this difference in recipes, Musken *et al.* (2008) used simulated intestinal fluids mimicking the ileum and colon to demonstrate differential expression of a major fimbrial subunit in sorbitol-fermenting EHEC while Polzin *et al.* (2013) found that EHEC proteins involved in nucleotide biosynthesis and the expression of Shiga toxins were increased in simulated ileal and colonic environments. In addition, it was found that outer membrane vesicle (OMV) production and cytotoxicity which is an important virulence factor as well as OMV-associated Shiga toxin 2a in EHEC was increased in both simulated ileal and colonic environments (75). On the other hand, OMV cytotoxicity and OMV-associated Shiga toxin 2a was not increased in DMEM which was confirmed with RT-qPCR for *stx_2a_* expression (75). These studies, in conjunction with the data presented here, highlight the importance of using a physiologically relevant environment when it comes to monitoring gene and protein expression in cells during gastrointestinal survival, colonization, and virulence. While we found that HG-DMEM is superior at inducing the expression of the virulence factor *ler*, we propose that SCF would be a useful medium to screen genes that have the potential to be involved in or required for colonization (Figure 5C-D).

### Concluding Statement

In this study, uncharacterized members of the *C. rodentium* Cpx regulon were investigated to determine if they were responsible for the avirulent phenotype exhibited *in vivo.* Our results demonstrate that neither *yebE, ygiB, bssR,* nor *htpX* are required for virulence, although they each require the Cpx ESR for maximal expression in multiple conditions and over different growth phases. In addition, we provide evidence that the dysregulation of virulence factors and fitness defects exacerbated in a simulated gut environment likely contribute to the colonization defect and attenuation of virulence exhibited by Δ*cpxRA* mutant cells.

## Acknowledgements

The authors thank the University of Alberta Molecular Biology facility for assisting with Sanger sequencing and Kat Pick for finding and suggesting the recipe of simulated colonic fluid used as well as for thoughtful discussion of results.

This research was supported by operating grants by the National Sciences and Engineering Research Council (NSERC) to TR and to SG.

## Tables

**Table S1. Strains and plasmids used in this study.**

**Table S2. Oligonucleotide primers used in this study.**

^1^All primers were designed in this study except for pNLP10_F/R (21)

^2^Restriction sites are underlined, enzymes indicated in primer name.

**Table S3. Average CPS/OD_600_ of three biological replicates 1-hour post-resuspension in LB and high-glucose DMEM with MOPS for cpxP-, yebE-, ygiB-, bssR-, and htpX-lux reporter plasmids.**

**^1^**Values (CPS/OD_600_) are the same as those represented in Figure 1A

**^2^***P-*value indicates significant difference between wild-type and Δ*cpxRA* mutant cells grown in the same condition (LB or HG-DMEM), Student’s *t-*test

## Figure Legends

**Figure S1. Luminescence on solid LB agar for reporters of confirmed Cpx regulon members and *cpxP-lux* in Cpx-regulated gene mutants.** (A) Wild-type and Δ*cpxRA* strains harbouring *lux*-reporter plasmids for each gene of interest and (B) wild-type and mutant strains harbouring *cpxP-lux* reporter plasmids were grown overnight, standardized to OD_600_ 1, serially diluted to 10^-6^, spotted on LB supplemented with kanamycin. Plates were pictured (top) after 18 hours of growth at 37°C and luminescence (bottom) was imaged using a ChemiDoc MP imaging system (Bio-Rad). All assays (A and B) were completed at least twice, with one representative experiment shown.

**Figure S2. Expression of *ygiB-* and *tolC-lux* over time indicates CpxRA dependent regulation.** (A-B) Wild-type (solid line) and Δ*cpxRA* (dotted line) cells harboring either *ygiB-lux* (blue) or *tolC-lux* (green) reporter plasmids and wild-type cells containing *cpxP-lux* (gray) were grown overnight then subcultured 1:100 in LB broth supplemented with kanamycin and grown to an OD of 0.4-0.6 at 37°C shaking. Two 1 ml aliquots of each subculture were spun down and resuspended in either (A) LB or (B) HG-DMEM without MOPS. The resuspended cells were added into a black walled 96-well plate and incubated at 37°C shaking. Luminescence was measured in counts per second (CPS) every hour for 6 hours starting at 0.5 hours post-resuspension. Reporter activity was plotted relative to OD_600_, and data represents the mean and standard deviation of three biological replicate cultures. (C) Diagram of *tolC* and *ygiBC* in the *C. rodentium* DBS100 genome. The CpxRA putative binding site is indicated with the black outlined box. Numbers in brackets indicate the number of nucleotides.

**Figure S3. Growth curve of luminescent reporter strains grown and measured over 12 hours.** Strains harboring *lux*-reporter plasmids were grown overnight and inoculated 1:100 in LB broth supplemented with kanamycin in a black walled 96-well plate and incubated at 37°C shaking. OD measurements were taken alongside luminescence measurements to ensure even growth between strains.

**Figure S4. CpxRA activity over time highlights differences in stresses experienced by cells based on culture conditions.** (A-B) Wild-type cells containing *cpxP-lux* were grown overnight then subcultured 1:100 in LB broth supplemented with kanamycin and grown to an OD of 0.4-0.6 at 37°C shaking. Two 1 ml aliquots of each subculture were spun down and resuspended in either (A) LB or (B) HG-DMEM buffered with 0.1M MOPS. The resuspended cells were added into two black walled 96-well plate and incubated at 37°C either shaking (dashed line) or statically (solid line). Plates were shook for 30 seconds followed by measuring the luminescence in counts per second (CPS) every hour for 6 hours starting at 0.5 hours post-resuspension. Reporter activity was plotted relative to OD_600_, and data represents the mean and standard deviation of three biological replicate cultures.

**Figure S5. Expression of the LEE master regulator, *ler,* over time in various conditions indicates reduced expression in the presence of CpxRA.** (A-D) Wild-type (solid line) and Δ*cpxRA* (dotted line) cells harboring *ler-lux* (pink) reporter plasmids and wild-type cells containing *cpxP-lux* (black) were grown overnight then subcultured 1:100 in LB broth supplemented with kanamycin and grown to an OD of 0.4-0.6 at 37°C shaking. Two 1 ml aliquots of each subculture were spun down and resuspended in either (A and C) low-glucose DMEM buffered with 0.1M MOPS or (B and D) HG-DMEM buffered with 0.1M MOPS. The resuspended cells were added into two black walled 96-well plate and incubated at 37°C either shaking or statically. Plates were shook for 30 seconds followed by measuring the luminescence in counts per second (CPS) every hour for 6 hours starting at 0.5 hours post-resuspension. Reporter activity was plotted relative to OD_600_, and data represents the mean and standard deviation of three biological replicate cultures.

